# HmtVar: a brand-new resource for human mitochondrial variations and pathogenicity data

**DOI:** 10.1101/355461

**Authors:** R. Preste, O. Vitale, R. Clima, M. Attimonelli

**Affiliations:** Department of Biosciences, Biotechnology and Biopharmaceutics, University of Bari, Bari 70126, Italy; Department of Medical and Surgical Sciences – DIMEC, Medical Genetics Unit, University of Bologna, Bologna 40126, Italy

## Abstract

Human mitochondrial data are currently of great interest for both clinicians and researchers, due to the involvement of mitochondria in a number of physiological and pathological processes. Thanks to new sequencing technologies and modern databases, the huge amount of information about mitochondrial genome variability can be exploited to gain interesting insights into the relationship between DNA variants, phenotypes and diseases. For this reason, we have developed the new HmtVar resource, a variant-focused database which allows to explore a dataset of over 30000 human mitochondrial variants together with their pathogenicity prediction. Mitochondrial variation data, initially gathered from the HmtDB platform, are further integrated with in-house pathogenicity assessments based on well-established variants pathogenicity evaluation criteria, as well as with a set of additional annotations from third-party resources. This approach led to a comprehensive collection of information of crucial importance for human mitochondrial variation studies and investigation of common and rare diseases in which the mitochondrion is involved to some extent.

HmtVar is accessible at https://www.hmtvar.uniba.it and its data can be retrieved using either a web interface through the Query page or a state-of-the-art API for programmatic access.

## Introduction

The mitochondrion is traditionally defined as the power-house of the eukaryotic cell and as such it has been considered by clinicians as involved in several pathologies, such as neurodegenerative diseases, diabetes, cancer and metabolic syndromes. However, the mitochondrion plays a pivotal role in many other biological processes, where it shows high variation in structure, proteomic composition and function differentiated in tissues and cell types. Hence the current great interest in mitochondria and disease, as confirmed by clinical literature^1,2,3^.

Recent advances in high-throughput sequencing techniques have provided an unprecedented amount of biological data, that is able to offer unevaluable insights into different life sciences questions. This acquires even more significance given the high number of biological sequences and related metadata that are available in public databases. Most of these data regards genomic variability, that can be exploited to achieve a better understanding of correlations between DNA variants, phenotypes and diseases.

Big public biological datasets can be used to assess human diversity. The large amount of information available allows to correlate phenotypic differences with genomic sequence data, in order to identify -with a certain degree of confidence- variants involved in determining some phenotypic traits. On the other side, some variations may simply represent neutral population polymorphisms, and can thus be flagged as non-pathogenic variants. This approach led to the development of several pathogenicity prediction tools, that are capable of estimating the “disease-causing value”, or pathogenicity score, of specific variants; this score is calculated through different algorithms that make use of the above-mentioned variability data.

Concerning the mitochondrial genome and its central role in a great number of diseases, during the last few years a huge quantity of online resources for mitochondrial data analysis has surfaced, with the common aim of producing a comprehensive knowledge about the onset and development of diseases in which mitochondria are involved. Some examples include the Mitomap^4^portal, the MSeqDR platform^5^and the HmtDB database^6^; however, a specific resource allowing to specifically query and retrieve mitochondrial variants with dedicated functional, structural, and population annotations and disease-related information is still lacking.

To this aim we have designed and implemented the resource HmtVar (https://www.hmtvar.uniba.it), a variant-centered database which is proposed as a comprehensive online platform that will allow clinicians and researchers to access human mitochondrial DNA variability and pathogenicity information. The high number of resources from which HmtVar’s data are integrated renders it one of the most exhaustive tools in this particular topic.

## Materials and methods

### Data sources

HmtVar is a variant-centered database which offers mtDNA variability and pathogenicity data (Fig. 1). The information available in HmtVar comes from different resources and spans several subjects in human mitochondrial genomics data. Variants data come from HmtDB^6^(https://www.hmtdb.uniba.it), given the high number of human mitochondrial genomes stored in it. The 39052 HmtVar variants are gathered from observed variations found in 34418 complete human mitochondrial genomes available on HmtDB, coming from either healthy and diseased subjects. Only complete genomes are here considered, i.e. those that do not lack any portion of the mitochondrial genome. Furthermore, additional potential variants integrate this dataset: these variants are not found in HmtDB genomes, but are determined based on possible substitutions for each given position in the mitochondrial genome. For this reason, while observed variants show a nucleotide variability score ranging from 0 to 1, for potential variants such value is equal to 0, as further detailed below.

**Figure 1.**
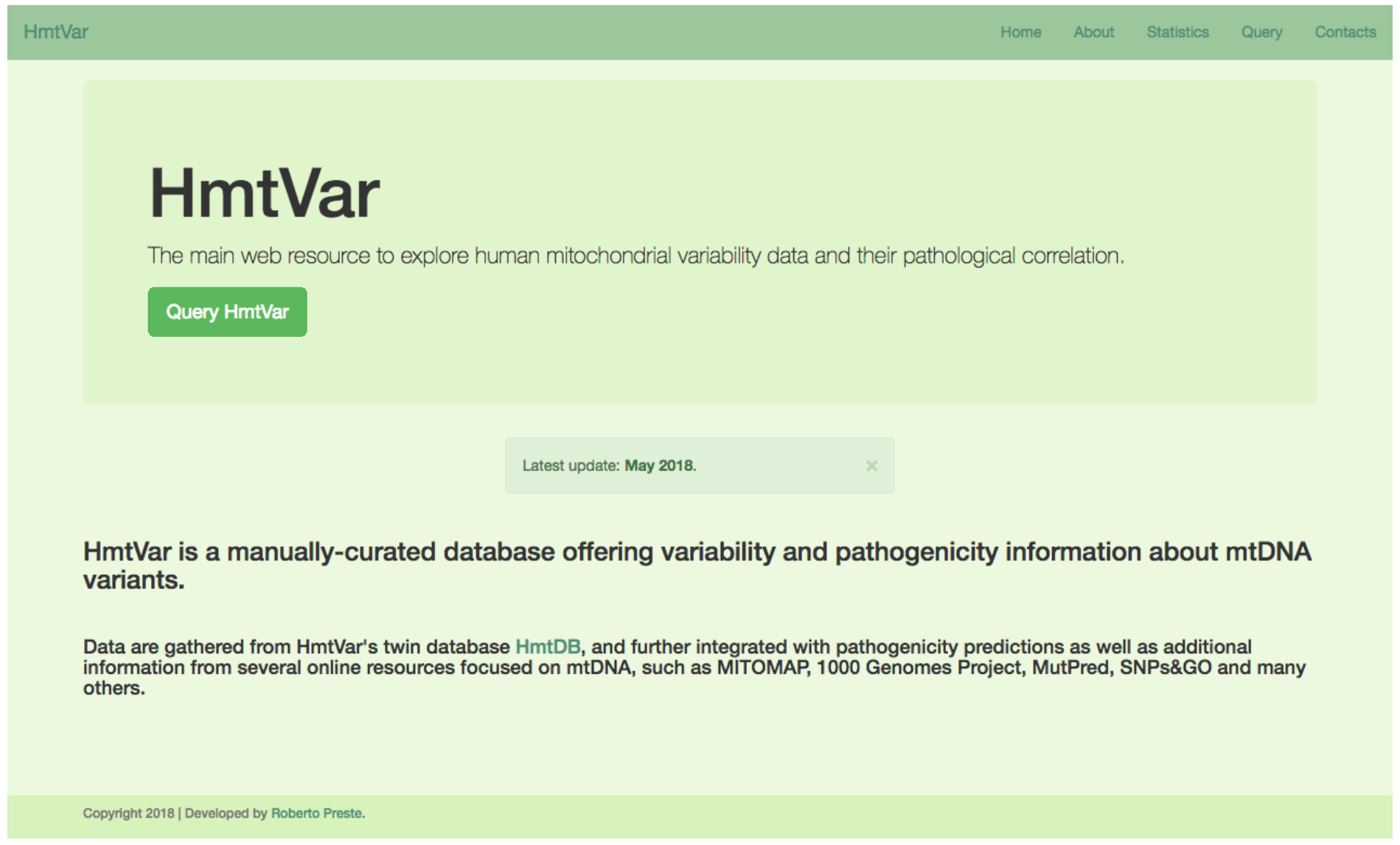
*HmtVar home page*

This allows the collection of one of the most extensive sets of mitochondrial variations, annotated with respect to the revised Cambridge Reference Sequence (rCRS, Accession Number NC_012920.1)^7^.

### Variants Site-Specific Variability

Site-specific variability calculations are performed by HmtDB, using the SiteVar algorithm for nucleotide variability^8^and the MitVarProt algorithm for aminoacid variability^9^; in both cases calculations are based on either healthy and diseased genomes, and are performed on the total number of genomes as well as on continent-specific subsets of them. The resulting variability score (ntVar) ranges from 0 to 1, with a higher value representing a lower functional constraint of the site. Alongside with variability data, allele frequencies are calculated for healthy, diseased and continent-specific genomes subsets; these data are also collected from HmtDB and stored in HmtVar.

### Variants Pathogenicity Estimation

#### Disease Scores Estimation

Disease Scores Estimations (DSE) for non-synonymous variants are based on the algorithm implemented in Santorsola et al^10^based on the weighted mean of the following six pathogenicity predictors: MutPred^11^, HumDiv- and HumVar-trained PolyPhen-2 models^12^, SNPs&GO^13,14^, PhD-SNP^15^and PANTHER^16^.

DSE for mt-tRNA variants, based on criteria established in Diroma et al^17^and Yarham et al^18^, have been improved by defining 4 additional macro-criteria, such as tRNA structure parameters, population frequency, heteroplasmy and functional studies evidence (Table 1). Within these macro-criteria, the following innovative points have been added:

a. Alterations of the cloverleaf-shaped secondary structure alteration determined by a nucleotide change, as already implemented in MitoTIP^19^;
b. Assessment based on post-transcriptional modifications;
c. Sites involved in 3D interactions;
d. Allele frequencies and further population data;
e. Contribution of the variant to a specific macro-haplogroup definition;
f. PhastCons^20^and PhyloP^21^conservation scores.

Moreover, corrective negative values have been established to support the benignity of variants. The total score ranges from 0 to 20, and is then normalized to a 0-1 range.

**Table 1.** *tRNA scoring system. Rows highlighted in blue are new properties introduced in the improved scoring system, while those highlighted in green represent criteria defined by Yarham et al and implemented by Diroma et al.*

| **tRNA Scoring Criteria** | **Yes** | **No** |
| --- | --- | --- |
| **tRNA structure parameters and reports** |  |  |
| Variant described as pathogenic by more than 1 report | 2 | 0 |
| Cloverfield-shaped 2D structure variation | 0.5 | 0 |
| Post-transcriptional modification | 1 | 0 |
| 3D interaction involved in folding | 1.5 | 0 |
| Variant described as benign by at least 1 report | -2 | 0 |
| **Frequency and population data** |  |  |
| PhastCons and PhyloP conservation | 1 | 0 |
| Variant frequency in patients > variant frequency in healthy | 1 | 0 |
| Variant frequency > 2% | -1 | 0 |
| Variant defining macro-haplogroup | -1 | 0 |
| 1000 Genomes data | -0.5 | 0 |
| **Heteroplasmy** |  |  |
| Heteroplasmy | 2 | 0 |
| **Functional studies evidence** |  |  |
| Molecular genetic analysis | 2 | 0 |
| Histochemical evidence | 2 | 0 |
| Biochemical defect in OXPHOS complexes | 2 | 0 |
| Cybrids/single fibre cells confirm pathogenicity | 5 | 0 |
| Cybrids/single fibre cells confirm benignity | -5 | 0 |

#### Disease Scores Threshold

Disease Scores Thresholds (DST) are defined by applying the bimodal distribution of disease score frequencies as described in Santorsola et al^10^. This procedure allowed to determine a value of 0.43 as non-synonymous variants’ DST, and a value of 0.35 as tRNA variants’ DST.

#### Nucleotide Variability Threshold

The nucleotide variability threshold (nt_var_T) values are defined by applying the empirical cumulative distribution^10^of nucleotide variability values for variants with a disease score above the DST. This procedure allowed to determine a nt_var_T of 0.0026 for non-synonymous variants and a nt_var_T of 0.0002315 for tRNA variants.

#### Tiers definition

In order to assign each non-synonymous or tRNA variant to a specific tier of pathogenicity, DST and nt_var_T have been considered, according to the following general rules:

**Table 2.** *General rules for variants’ pathogenicity assignment*.

| **Tier** | **Disease Score range** | **Nucleotide Variability range** |
| --- | --- | --- |
| Polymorphic | $DS < DST$ | $nt\_var > nt\_var\_T$ |
| Likely Polymorphic | $DS < DST$ | $nt\_var \leq nt\_var\_T$ |
| Likely Pathogenic | $DS \geq DST$ | $nt\_var > nt\_var\_T$ |
| Pathogenic | $DS \geq DST$ | $nt\_var \leq nt\_var\_T$ |

Taking advantage of the specific thresholds defined above, the final pathogenicity tiers have been defined as detailed in Table 3.

**Table 3.** *Non-synonymous and tRNA variants’ specific rules for pathogenicity assignment*.

| Tiers | Disease Score range | Nucleotide Variability range |
| --- | --- | --- |
| **Non-synonymous variants** |  |  |
| Polymorphic | DS < 0.43 | nt_var > 0.003282 |
| Likely Polymorphic | DS < 0.43 | nt_var ≤ 0.003282 |
| Likely Pathogenic | DS ≥ 0.43 | nt_var > 0.003282 |
| Pathogenic | DS ≥ 0.43 | nt_var ≤ 0.0026 |
| **tRNA variants** |  |  |
| Polymorphic | DS < 0.35 | nt_var > 0.0002315 |
| Likely Polymorphic | DS < 0.35 | nt_var ≤ 0.0002315 |
| Likely Pathogenic | DS ≥ 0.35 | nt_var > 0.0002315 |
| Pathogenic | DS ≥ 0.35 | nt_var ≤ 0.0002315 |

### Implementation

HmtVar is built upon the Python Flask framework^22^and uses the SQLite^23^database to manage its data. The web interface was developed using Bootstrap^24^, and the update procedure takes advantage of the Nextflow pipeline management system^25^.

The choice of implementing HmtVar’s back-end functionality using Python Flask was made out of the need for a lightweight yet efficient framework for both building a reliable web service and to retrieve and parse information from the underlying database. Python Flask integrates very well with SQLite databases and offers a set of tools to perform database construction, querying, editing and versioning right out of the box. SQLite, on the other hand, allows for a simpler and faster data storage and retrieval, with respect to legacy database engines; this is fundamental given the high amount of genomic information collected.

In addition, it was possible to deploy a comprehensive Application Programming Interface (API) without relying on other services, such that researchers and developers can access HmtVar’s data in a programmatic way and integrate them into their applications.

HmtVar was developed with one of the key ideas being ease of access for everyone and from every device, thus the Bootstrap library was employed for developing its front-end. As such, HmtVar can be accessed from either desktop and mobile devices without any loss in user experience.

One of the aims of HmtVar is to provide users with data which are always up-to-date. This means collecting new variant entries and variability data as soon as they are available in HmtDB, as well as gathering additional information from several third-party resources. Specific software to retrieve new available information from each data source was first developed; this set of scripts was then aggregated using the Nextflow pipeline manager, which is capable of parallelizing the different retrieval and parsing tasks, thus considerably speeding up the entire updating process.

## Results and discussion

### Data statistics

HmtVar currently hosts a grand total of 39052 human mitochondrial variants, classified as observed or potential variants, and distinguished as follows (Fig. 2):

- 33622 non-synonymous variants located in protein coding genes (CDS);
- 4483 variants located in tRNA genes;
- 563 variants located in the D-Loop;
- 384 variants located in rRNA genes (RNR1 and RNR2).

**Figure 2.**
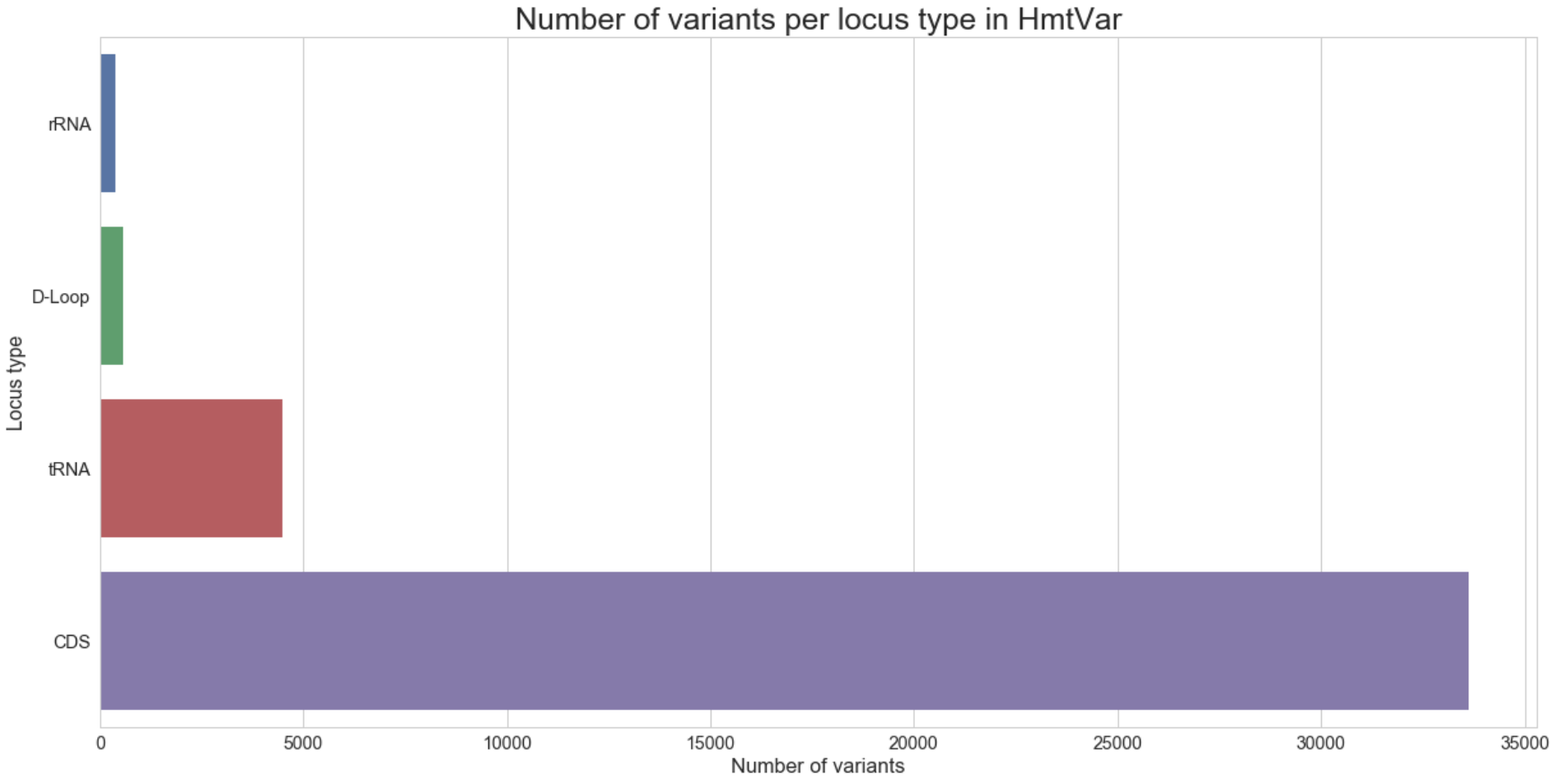
*Variants in HmtVar per locus type*

Variants located in the D-Loop and in rRNA genes only represent a subset of the total number of potential variants belonging to these regions, and will be integrated with further HmtVar updates.

Each variant has been assigned to one of the four tiers of pathogenicity (polymorphic, likely polymorphic, likely pathogenic, pathogenic), as already described in Table 2.

Pathogenicity predictions are offered for 38105 of the total number of variants hosted on HmtVar, allowing to classify them as pathogenic (17172), likely pathogenic (1562), polymorphic (2372) and likely polymorphic (6814) (Fig. 3A).

**Figure 3.**
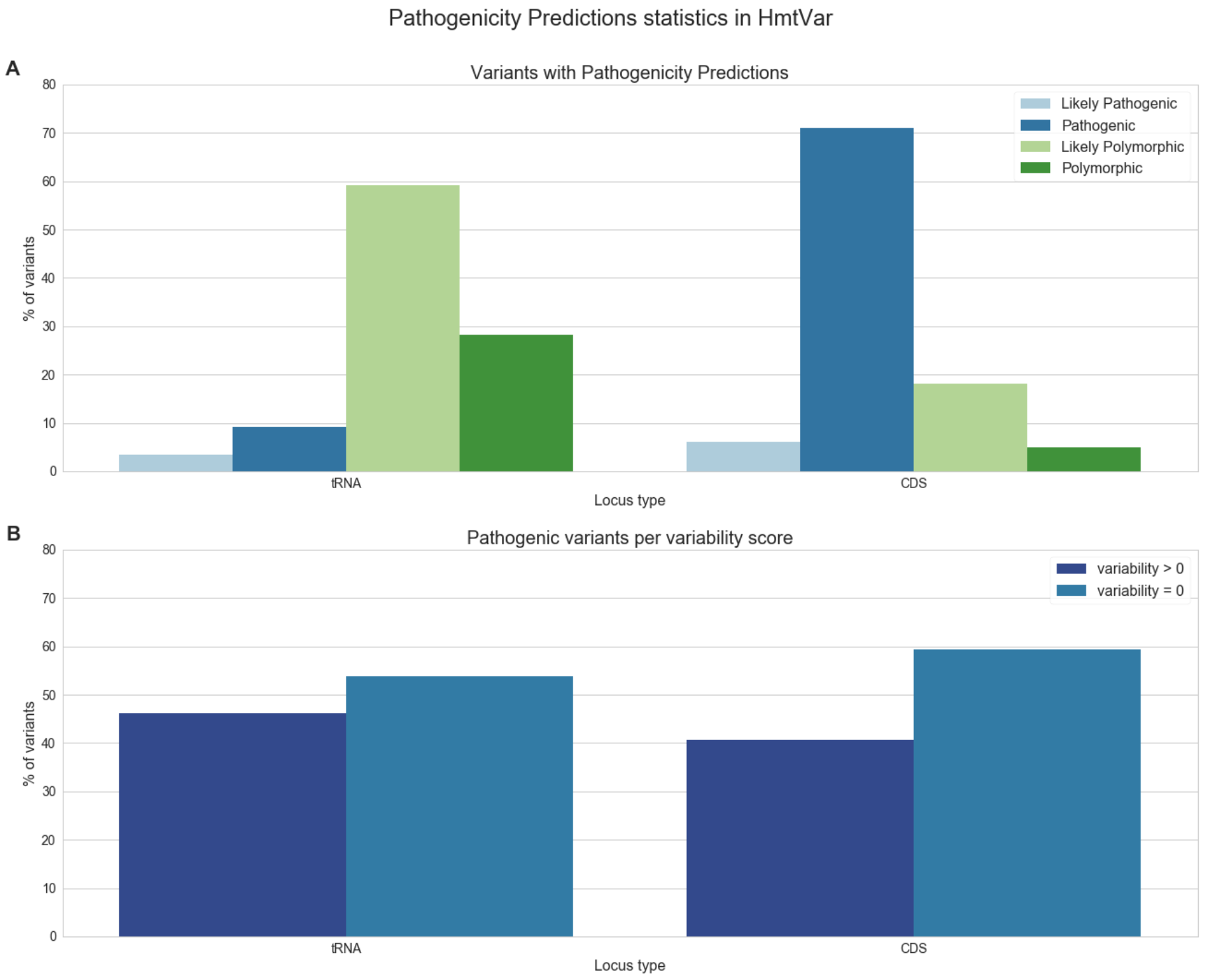
*A: Percentage of variants with defined pathogenicity predictions. B: Percentage of variants classified as pathogenic distinguished as observed (variability > 0) and potential (variability = 0)*.

For both tRNA and protein coding genes, most pathogenic variants are potential, with pathogenic tRNA potential variants being 59* of the total number of tRNA variants, and pathogenic non-synonymous potential variants in protein coding genes being 53* of their total number (Fig. 3B).

### Interface

HmtVar is accessible at https://www.hmtvar.uniba.it/, and offers either a web interface and a RESTful API to query its content. Using the Query web page, data can be queried using several search parameters, from more broader criteria (i.e. variants located in a certain locus or with a particular pathogenicity prediction) to more definite ones, such as specific variant position or variability value.

Each variant’s annotation is shown in a Variant Card, which gathers all the information available for the selected variant and arranges them in a neat way using different tabs (Fig. 4).

**Figure 4.**
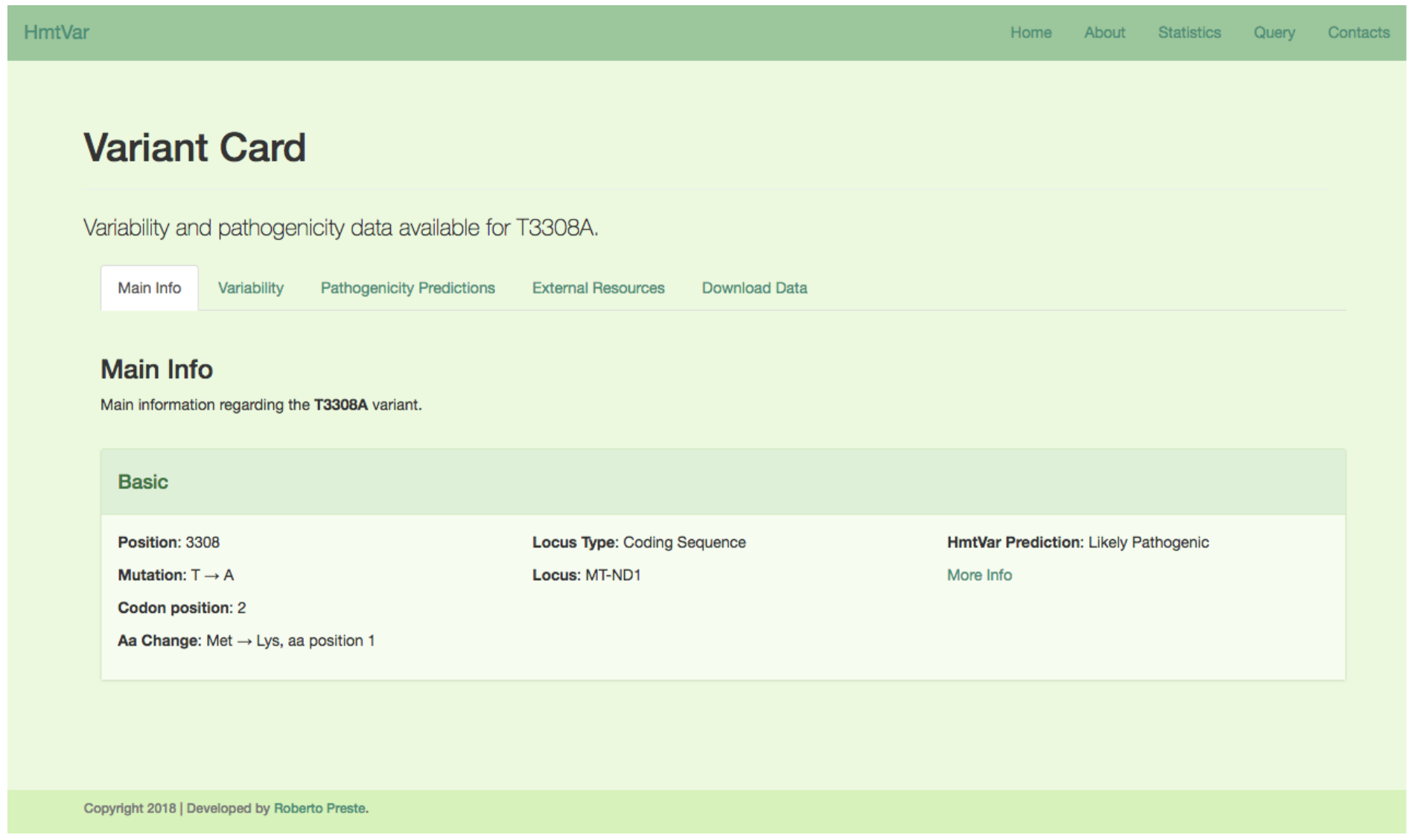
*HmtVar Variant Card*

The **Main Info** tab reports the variant’s basic information, like its location, the consequent aminoacidic change or tRNA annotations (if applicable), haplogroups and macro-haplogroups defined by the variant, as well as HmtVar’s pathogenicity prediction.

The **Variability**tab shows nucleotidic and aminoacidic variability values, for both healthy and diseased genomes; in the same tab, healthy, diseased and continent-specific allele frequencies are also reported.

The **Pathogenicity Predictions**tab shows a more detailed view of HmtVar’s pathogenicity prediction together with the related disease score; in addition, some more pathogenicity predictions and scores calculated by external online tools such as MutPred^11^, PolyPhen-2^12^, PANTHER^16^, PhD-SNP^15^and SNPs&GO^13,14^are shown in specific panels. When these data are available, both the qualitative pathogenicity classification and the quantitative pathogenicity prediction score are reported, as calculated by each of these resources.

In the **External Resources** tab, a set of additional variant information is shown; these data come from many online resources, ranging from clinical significance (ClinVar^26^, OMIM^27^) to variant description (dbSNP^28^and Mamit-tRNA^29^), to variant homo/heteroplasmy as reported by Mitomap^4^and 1000Genomes^30,31^datasets. Links to the original sources of information are always provided for consistency.

Finally, the **Download Data** tab allows users to download each variant’s data for offline use. Data are provided in a JSON-formatted file, which contains all the available information for the selected variant, distinguished based on the above-mentioned categories.

### API

In addition to the Query page functionality, HmtVar allows to retrieve variants using a dedicated Application Programming Interface (API), which is most suited for users needing to access and download data in a programmatic manner. Valid API calls will return one or more results formatted as a JSON string, for an easy parsing of information.

Requests to the HmtVar API can be made to https://www.hmtvar.uniba.it/api/main/, and can accept one of the following arguments:

- position/<nt_pos> to retrieve variants located in a given position in the human mitochondrial genome; the <nt_pos> parameter can also accept a list of positions (separated by commas) or a range of positions (separated by a dash);
- mutation/<mut> to retrieve one or more specific variants, identified by reference allele, position and alternate allele; the <mut> parameter can be formatted in one of the following ways:

- [ref][pos][alt] to query for a specific variant with the given reference allele ([ref]), position ([pos]) and alternate allele ([alt]);
- [pos][alt] to query for a specific variant with the given position ([pos]) and alternate allele ([alt]);
- [ref][pos] to query for all available variants starting with the reference allele ([ref]) and position ([pos]);
- locus/<loc> to retrieve all variants located in the given mitochondrial locus;
- pathogenicity/<patho> to retrieve all variants for which a specific pathogenicity prediction is available (accepted values for the <patho> argument include pathogenic, polymorphic, likely_pathogenic, likely_polymorphic).

When returning a single variant, the API will provide the complete set of available information about that specific variant, exactly like the data shown in the Download Data tab of a VariantCard.

When returning a list of variants, instead, each variant’s entry will report the URL to directly access that variant’s complete data, as well as a limited set of basic information about that variant.

### RD-Connect compliant API

RD-Connect is a comprehensive platform that integrates databases, patient registries, data analysis tools and biobanks for rare disease research^32^. In order to collect and integrate data from a broad range of bioinformatics resources, RD-Connect established a common API that data providers can adopt; using a set of standardized arguments for API calls, RD-Connect is then able to retrieve, parse, integrate and redistribute third-party data. To further distribute mitochondrial variants information, HmtVar also offers a second form of API in addition to the one previously described, created to be compliant with the RD-Connect platform specifications.

Data coming from both HmtDB and HmtVar are particularly useful for RD-Connect to estimate mitochondrial variants pathogenicity and variability, thus we developed a compliant API based on RD-Connect’s common API specifications. Variants data can be retrieved with a GET request to https://www.hmtvar.uniba.it/rdconnect? using one or more of the following arguments to search for specific data, concatenating them with &:

a. gene_symbol=<string> to retrieve variants located in the given mitochondrial gene;
b. gene_id=<string> to retrieve variants located in the given mitochondrial gene id;
c. variant_start=<int> to retrieve variants starting on the given mitochondrial position;
d. variant_end=<int> to retrieve variants involving more than a single nucleotide and ending on the given mitochondrial position;
e. variant_referenceBases=<char> to retrieve variants for which the reference nucleotide is the given nucleotide;
f. variant_alternateBases=<char> to retrieve variants with the given alternate allele.

Requests to this API will return a JSON-formatted string containing a “success” key, whose value can be either “true” if the requested data are available on HmtVar or “false” otherwise, and a “url” key which reports the permanent link to the corresponding HmtVar VariantCard. This link can then be exploited by RD-Connect to collect and integrate HmtVar variants data into their platform.

## Conclusions

HmtVar offers a wide range of information regarding human mitochondrial genome variants, representing one of the most comprehensive resources for genomic variation studies. The broad set of mitochondrial data hosted on several different online sources was exploited to build a unique aggregated data repository which will be able to fulfil the clinicians needs when looking for pathogenicity information about mitochondrial variants.

Pathogenicity predictions for variants located in mitochondrial tRNA and coding genes allow to assess each variation’s deleteriousness based on a pathogenicity consensus obtained from different tools, thus offering a certain degree of reliability.

These data can be accessed via a web page as well as an API, with the latter option allowing researchers and developers to integrate information available on HmtVar into their custom applications.

Future implementations will focus on extending HmtVar’s dataset beyond observed variants to embrace the whole set of potential variations with respect to the rCRS reference sequence; in addition, calculations of pathogenicity prediction for rRNA and D-Loop variants will be performed, in order to offer a full overview of human mitochondrial variability and pathogenicity.

## Acknowledgment

The RD-Connect compliant API for HmtVar was developed in the context of the 2017 ELIXIR Implementation Study “Integration of the ELIXIR-IIB HmtDB resource into RD-Connect”.

